# Assessing sustainability in North America’s ecosystems using criticality and information theory

**DOI:** 10.1101/330415

**Authors:** Elvia Ramírez-Carrillo, Oliver López-Corona, Juan C. Toledo-Roy, Jon C. Lovett, Fernando de León-González, Luis Osorio-Olvera, Julian Equihua, Everardo Robredo, Alejandro Frank, Rodolfo Dirzo, Vanessa Perez-Cirera

**Affiliations:** Doctorado en Ciencias Agropecuarias, Universidad Autónoma Metropolitana-Xochimilco, Ciudad de México, México; Cátedra CONACyT, Comisión Nacional para el Conocimiento y Uso de la Biodiversidad (CONABIO), México; Red Ambiente y Sustentabilidad, Instituto de Ecología A.C. de México (INECOL); Centro de Ciencias de la Complejidad (C3), Universidad Nacional Autónoma de México, Ciudad de México, México; Instituto de Ciencias Nucleares, Universidad Nacional Autónoma de México, Ciudad de México, México; School of Geography, University of Leeds, Leeds, LS2 9JT, United Kingdom; Royal Botanic Gardens, Kew, Richmond, Surrey TW9 3AB; Departamento de Produccion Agricola y Animal, Universidad Autónoma Metropolitana-Xochimilco, Ciudad de México, México; Posgrado en Ciencias Biológicas, Faultad de Ciencias, Universidad Nacional Autónoma de México; Comision Nacional para el Conocimiento y Uso de la Biodiversidad, México; Miembro del Colegio Nacional; Biology Department, Stanford University, US; Instituto de Investigatión para el Desarrollo con Equidad (EQUIDE), Universidad Iberoamericana, Ciudad de México, México

## Abstract

Sustainability is a key concept in economic and policy debates. Nevertheless, it is usually treated only in a qualitative way and has eluded quantitative analysis. Here, we propose a sustainability index based on the premise that sustainable systems do not lose or gain Fisher Information over time. We test this approach using time series data from the AmeriFlux network that measures ecosystem respiration, water and energy fluxes in order to elucidate two key sustainability features: ecosystem health and stability. A novel definition of ecosystem health is developed based on the concept of criticality, which implies that if a system’s fluctuations are scale invariant then the system is in a balance between robustness and adaptability. We define ecosystem stability by taking an information theory approach that measures its entropy and Fisher information. Analysis of the Ameriflux consortium big data set of ecosystem respiration time series is contrasted with land condition data. In general we find a good agreement between the sustainability index and land condition data. However, we acknowledge that the results are a preliminary test of the approach and further verification will require a multi-signal analysis. For example, high values of the sustainability index for some croplands are counter-intuitive and we interpret these results as ecosystems maintained in artificial health due to continuous human-induced inflows of matter and energy in the form of soil nutrients and control of competition, pests and disease.

## Introduction

Sustainability has been defined in many ways, but the most frequently-quoted definition is from *Our Common Future*, also known as the Brundtland Report: ‘‘Sustainable development is development that meets the needs of the present without compromising the ability of future generations to meet their own needs”. The vagueness of this statement may suit the need for flexibility in policy objectives, but attempts to bring greater precision to implementation have long been thwarted by multiple possible interpretations [1]. So, despite considerable interest in the core idea of sustainability [2, 3], it remains a poorly demarcated concept, eluding mathematical definition except within the scope of a few restricted disciplines, and excluding fundamental laws such as entropy [4]. There is no widely-accepted, precise, and testable multidisciplinary definition of sustainability; and perhaps more importantly there is no general theory of the subject, thereby preventing rigorous analysis and evidence-based policy formulation.

Without precise mathematical definitions, a general theory, and a testable hypothesis, it is virtually impossible to apply the scientific method to make progress in any area of study. For example, together with economics and social, ecology is one of the three dimensions of sustainability, yet macroecology is woefully underrepresented in sustainability science [5]. While ecological principles are central to sustainability, their use has been largely concerned with socio-environmental interactions, such as physical limits on resource use by energy-demanding technology-based human societies. For example, Burger et al. (2012) describe calculations that are consistent with analyses reporting peak oil, fresh water, and phosphate, to examine how global stocks of these important resources affect the patterns of global consumption decline and the likelihood of global depletion [6, 7].

From a basic ecological perspective, sustainability encompasses “the ability of one or more entities, either individually or collectively, to exist and thrive (either unchanged or in evolved forms) for lengthy timeframes, in such a manner that the existence and flourishing of other collectivities of entities is permitted at related levels and in related systems” [8]. From the multiplicity of elements considered in this definition, we identify two core aspects relevant to ecological sustainability: ecosystem stability and health [9]. As a whole, the concept of sustainability faces important challenges if it is to consolidate as a scientific discipline. In this paper, we focus on the ecological dimension of sustainability, particularly stability and health, using a thermodynamic-informational framework. When we discuss stability we are doing so in a statistical sense rather that in a formal system dynamics way, calculating for example, Routh-Hurwitz conditions. We consider that this approach provides precise mathematical concepts and takes into account fundamental constraints in ecology and system dynamics.

### Ecosystem Health

Ecosystem health is a diffuse concept that has been defined several times since the late 1980’s [10]. This conceptual diversity has given rise to different measurement methods, which in turn have generated a wide range of narratives related to ecosystem health [11]. Ultimately it has become an ongoing priority for governments, scientists and managers around the world [5].

Originally, ecosystem health was conceived within a control-optimization perspective in which health is defined as a desired management target or reference condition [12, 13].

ure 1 lays out a schematic relationship between the ontologies that result from different perspectives. The top of the figure represents the ‘Natural’ perspective for which ecosystem health is usually defined through structure assessment and ecological functions. This involves the measurement of certain indicators, such as the lack of algal blooms in rivers [14, 15]; food web performance [16]; nutrient recycling and maintenance of biodiversity (US National Research Council, 2005), and resilience to external perturbations [17]. The bottom of the figure represents the ‘Human’ perspective that understands the issue from a managerial standpoint, focusing on optimization and control.

The definition of ecosystem health has entered the realm of multi- and interdisciplinarity (horizontal line in Fig. 1). At the level of multidisciplinarity, ecosystem health can be characterized by ecosystem services such as the provision of clean drinking water [18]. As the idea of sustainability has permeated society, ecosystem health has become increasingly associated with the integration of environmental, economic and human domains [10, 19].

**Fig 1.**
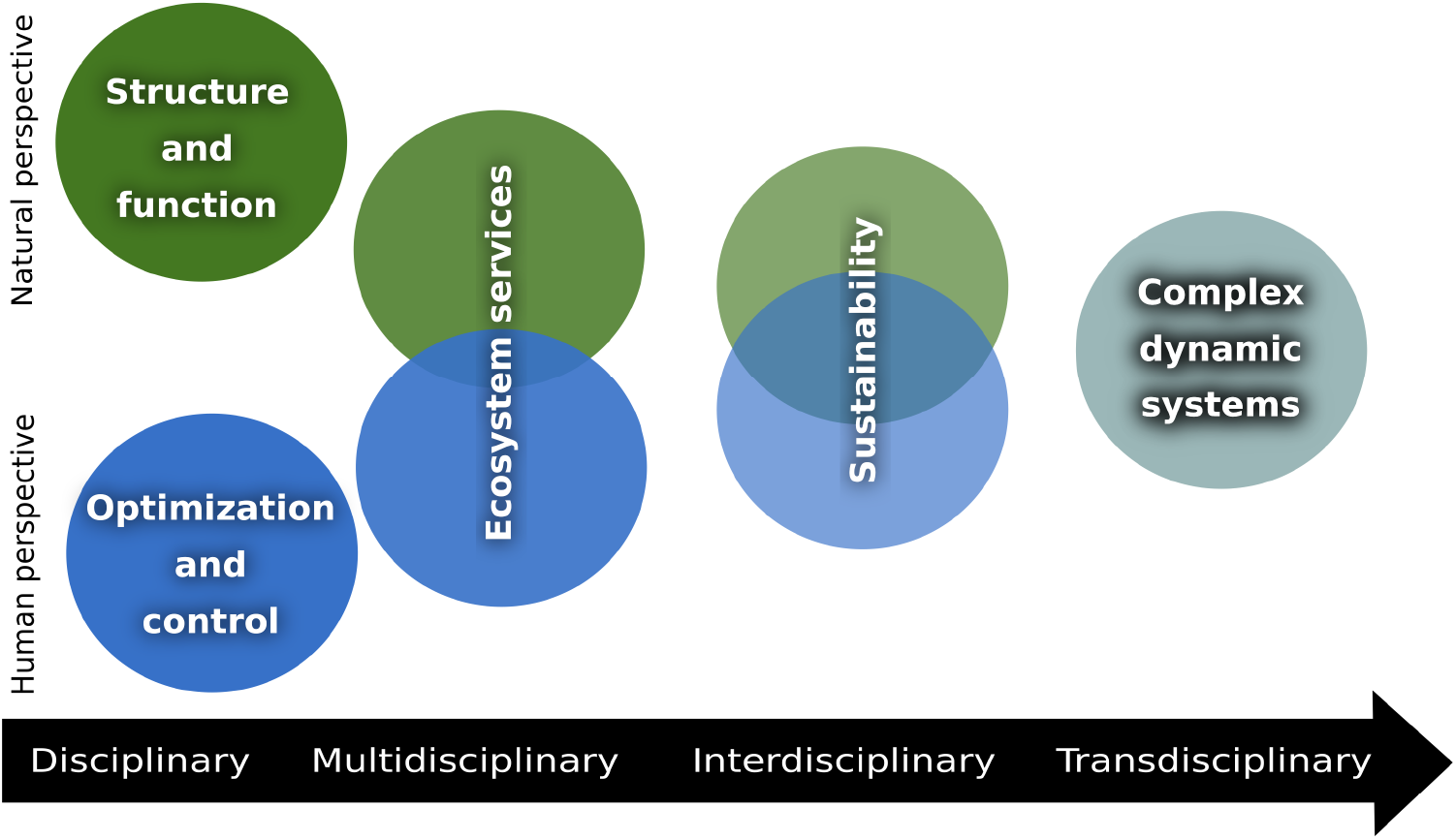
The narrative surrounding ecosystem health has shifted from a very disciplinary framework, through multi, inter and transdisciplinarity, before finally being defined in terms of complex systems.

Finally, the shift towards interdisciplinarity in science has led to a complexity-based approach in which ecosystem health is conceived as a property of a complex system. A dynamic system is characterized by a set of (state) variables. When assigned a particular set of numeric values, these variables define the state of the system. There are also evolution rules that describe the way in which the system transitions from any one state to any other. Although the very definition of a complex system is still under active discussion [20–24], in general a complex system emerges from a sufficiently large number of elements that have strong enough (usually non-linear) interactions, or when the state space changes fast enough in terms of observer’s scales of observation. These qualities make it impossible to describe the behavior of the system in terms of the simpler behavior of its components.

Within this narrative of system dynamics, complexity is usually studied by analyzing the time series of the fluctuations in state variables that have been identified as central to the dynamics of the system [25]. Ultimately, they are at the center of the modern description of out-of-equilibrium dynamics [26].

A typical analytical method is studying the time series through spectral and fractal analysis, in particular through the Power Spectral Density (PSD) or Detrended Fluctuation Analysis (DFA). It is often the case that these fluctuations exhibit scale invariance (e.g. when the power spectrum follows a power-law (*S* ~ *f^−β^*), in which case it is customary to compare and classify fluctuation dynamics according to their similarity to three archetypal classes of noises: white (*β* ~ 0), pink (*β* ~ 1) and Brownian (*β* ~ 2) [27–30].

It has also been reported in the literature that several complex systems display behaviors related to dynamic criticality, usually associated with some kind of scale invariance, and in many cases with pink noise [27, 28, 31, 32].

Following this line of thought, several authors have found evidence of dynamic criticality in physiological processes such as heart activity, and have posited that it may be a key feature of a healthy state. [33–35]. Some authors [36] strongly relate healthy hearts associate scale-invariant noise in the region around 1/f noise and provide medical evidence for it. In a recent paper reviewing criticality in the brain, [37] state that i) Criticality is a widespread phenomenon in natural systems that provides a unifying framework that can be used to model and understand brain activity and cognitive function, and ii) that there is substantial evidence now supporting the hypothesis that the brain operates near criticality.

Nevertheless, from a theoretical standpoint, the universality of criticality is still under examination and is known as the Criticality Hypothesis, which states that systems in a dynamic regime shifting between order and disorder, attain the highest level of computational capabilities and achieve an optimal trade-off between robustness and flexibility. Recent results in cell and evolutionary biology, neuroscience and computer science have great interest in the criticality hypothesis, emphasizing its role as a viable candidate general law in the realm of adaptive complex systems (see [38] and references therein).

Our proposal in this paper, to address ecosystem health, is based on the criticality framework and we measure it as the combination of scale invariance (as power laws in Power Spectra) and a balance between adaptability and robustness (dynamic in the neighborhood of a 1/f noise type). In this regard, [39] have pointed out that “the very existence of such ubiquitous power laws implies the existence of powerful constraints at every level of biological organization. The self-similar power law scaling implies the existence of average, idealized biological systems, which represent a 0th order baseline or point of departure for understanding the variation among real biological systems. Real organisms can be viewed as variations on, or perturbations from, these idealized norms due to influences of stochastic factors, environmental conditions or evolutionary histories”. This scale invariance property manifests itself, for example, as power law behavior. These power laws appear in countless phenomena including the statistics of earthquakes, solar flares, epidemic outbreaks,etc. [40–43]. They are also a common theme in biology [36, 44–47]. Of particular interest for this paper are the examples of many physiological and clinical time-series data that have a spectrum that decays as a power of the frequency. This effect is often called 1/f noise, although powers of the frequency, f, may appear [48]. Also, patterns of human and animal mobility often exhibit scale-free features [49–52]. Moreover, a number of commonly observed statistical patterns of natural-world data –such as Zipf’s law [42, 53–55], Bendford’s law [56, 57], and Taylor’s law [58, 59] - stem from underlying scale invariance, i.e. power-law distributions [60].

This universality of power laws may be due, as [61] proposes, as a result of the optimization of energy, matter and information transport. The proposed common mechanism underlies the idea that living things are sustained by the transport of materials through linear networks that branch out to supply all parts of the organism [61] and involves three principles or assumptions. First, in order for the network to supply the entire volume of the organism, a space-filling fractal-like branching pattern is required. Second, the final branch of the network (such as the capillary in the circulatory system) is a size-invariant unit. And third, the energy required to distribute resources is minimized; this final restriction is basically equivalent to minimizing the total hydrodynamic resistance of the system. The authors then claim that scaling laws arise from the interplay between physical and geometric constraints implicit in these three principles.

Furthermore, [62] showed that scale invariance emerge only at critical temperature levels of a two dimensional Ising model – temperature levels in which the correlation length diverges, which in practice means that the correlation length becomes very large when compared to the scales of interaction of the system. And they also support the conclusion that this property may be the key to the robustness and adaptability of complex systems.

A balance between robustness and adaptability has already been recognized as an important feature of sustainability by [63]. The authors state that sustainable systems tend to be in an optimal regime where the capacity for the system to undergo evolutionary change or self-organization consists of two aspects: i) It must be capable of exercising sufficient directed power (ascendency for them, robustness for us) to maintain its integrity over time and, on the other hand, ii) it must simultaneously possess a reserve of flexible actions (adaptability in our narrative) that can be used to meet the exigencies of novel disturbances. Then the authors argue that “systems with either vanishingly small ascendency or insignificant reserves are destined to perish before long. A system lacking ascendency has neither the extent of activity nor the internal organization needed to survive. By contrast, systems that are so tightly constrained and honed to a particular environment appear “brittle” in the sense of Holling (1986) or “senescent” in the sense of Salthe (1993) and are prone to collapse in the face of even minor novel disturbances. Systems that endure – that is, are sustainable – lie somewhere between these extremes”. In our case, that optimal regime that lies in-between is the criticality, mainly characterized by scale invariance.

If we study a system by its time series, and it is well accepted that this must be done on the time-series of the fluctuations instead of the original state variable, then a traditional place for looking for scale invariance is in its power spectrum *S* ~ *f^−β^*. When this happens, as the autocorrelation functions as the inverse Fourier transform of the power spectrum of the signal *C*(*τ*) = *F*^−1^(*S*), then applying a scale transformation in the time domain, *τ* → *τ*′ = *aτ*, we obtain the autocorrelation function of the type

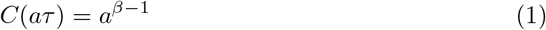

for which the general solution of the equation is also a power law. In this way, the correlations are zero for white noise, large for brown noise, and then pink noise is between no correlation (that we associate with adaptation) and high correlation (that we associate with robustness). Under this narrative, we propose that criticality is recognizable in coarse grain by power laws in the power spectrum, and then the system will be more critical if it is in the vicinity of *beta* = 1 (i.e., pink noise). When we relate criticality with health, our proposal is in good agreement with [64], who found that scale invariant and 1/f noise satisfy the unifying concept that physiological complexity (pink is more complex that white or brown noise) is fundamentally related to the adaptive capacity of the organism, which requires integrative, multiscale functionality. In contrast, disease states, as well as aging, may be defined by a sustained breakdown of long-range correlations.

Although we recognize that more work is needed to anchor the theory of criticality to health in general, or to ecosystem health in particular, we nevertheless consider that there is enough empirical evidence of the former, and there are justified reasons to believe it could be valuable in the future.

Following this line of thought, several authors have found evidence of this dynamic criticality in physiological process such as heart activity, and have speculated that it may be a key feature of a healthy state. [33–35]

Our proposal in this paper for measuring ecosystem health (Fig. 2) is based on this idea of dynamic criticality as the combination of scale invariance and balance between adaptability and robustness.

**Fig 2.**
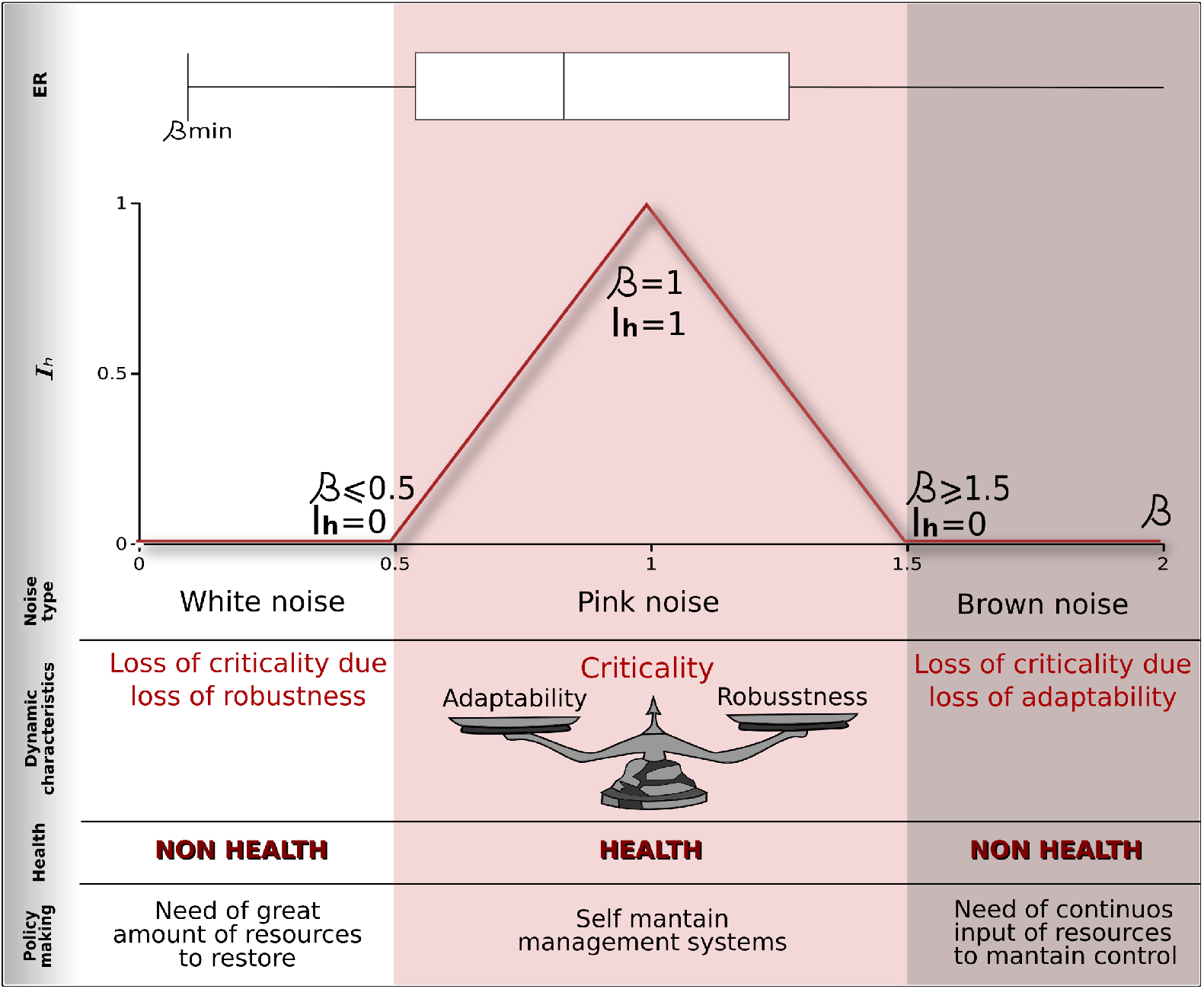
Our proposal for measuring the dynamic dimension of ecosystemic health is based on the idea of criticality as the combination of scale invariance and balance between adaptability and robustness (pink noise). By combining a scale invariance index based on BIC values with the value of the scalar coefficients (beta) in power spectra, we propose an Ecosystemic Health Index, whose maximum for beta values equals 1, and that is associated with a balance between adaptability and robustness. In this way, an ecosystem may lose health by losing robustness and exhibiting white-noise dynamics, or by losing adaptability leading to Brownian-noise dynamics.

Initially we asked which environmental signal could be used as an analog of a systemic physiological variable such as heart rate, and that is one of the most widely used experimental variables?

We consider that potential candidates should be related to soil because it is a complex system [65] that integrates several scales as well as the main ecosystemic processes. On the one hand, its dynamics are defined by the interaction of different subsystems such as the biosphere, atmosphere, geosphere and hydrosphere and all their components [66], which in turn interact in the geographic space, generating different pedogenetic processes related to climate or geoforms [67–69]. On the other hand, according to [65], carbon flows also connect ecosystems in the temporal dimension, since carbon persists in the soil as a kind of biogeochemical memory [70].

Following the above logic, and since soil plays a central role in the flux of CO_2_, we decided to analyze the Ameriflux database that measures energy and matter (mainly CO_2_) fluxes in America (mostly North America).

### Ecosystem Stability (Out-of-equilibrium Thermodynamics)

Following [71], let us consider an abstract resource space in which we can define a vector 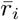 that represents a particular socio-ecological entity (a species, an environmental entity such as a wetland, or a human community). In this way, the projection of the entity’s vector 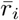 over a resource axis represents the root mean square value of the resource gradient that the entity *i* requires to subsist.

[71], proposes that the movement of an entity in this space implies changing the resource gradient requirements as well as the strength of the interactions with the other entities. Thus, an entity vector 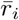 may exhibit length changes in the form of vibrations (i.e. changes in a species population), small direction changes in the form of rotations (i.e. niche plasticity) and larger changes in its direction (i.e. niche evolution).

Helmholtz discovered that, when left alone, all systems tend to more stable states of greater longevity by reducing their free energy *F* as defined by

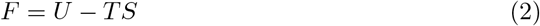

where *U* is the internal energy of the system given by interactions between the internal constituents, *T* is its temperature (associated with randomness) and *S* its entropy. Then, given the restrictions of the resource space, this optimization problem guides the dynamics of the exploration of this space and hence the evolution of the strengths of the interactions between entities and between entities and environment. Stability is reached by minimizing Helmholtz free energy, and one way to achieve this is to maximize the entropy of the whole system (not just a single component) [72].

### Ecosystem Stability (Fisher Information)

Mayer and co-workers [3] have proposed that Fisher information offers a robust method to assess the stability of a system over time, being essentially able to aggregate multiple variables, each one capturing different aspects of a system, and outputting a global indicator of stability.

Following [73] and [3] let us consider the basic problem of estimating the real value of a state variable *θ*. The estimation comes from an inference process from imperfect observation *y* = *θ* + *x* in the presence of some random noise *x*.

This kind of measurement-inference process will hence be called “smart measurement” of *θ* whose result is an estimator 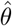 that is function of imperfect observation 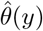.

This is a closed system, meaning that it’s well described by 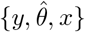 without the need to consider additional sources of noise. Consider also that the estimator is unbiased in terms of being a good estimator on average 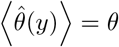. In this case, the mean-square error obeys the Cramer-Rao inequality

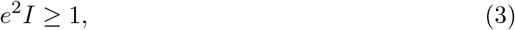

where *I* is the Fisher Information of the system, calculated as

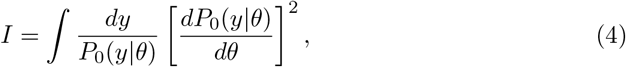

in which *P*_0_ (*y*|*θ*) is the probability density function of measuring a particular value of *y* given the true value *θ* of the state variable in question. Then, since the error decreases as information increases, Fisher information may be understood as the quality of the estimation *θ* from a smart measurement.

Then if the system is characterized by a phase space with m state variables x¿ that define the phase vector *s* = (*x*_1_, …, *x_i_*, …, *x_m_*) associated with a smart measurement y, then we can prove that

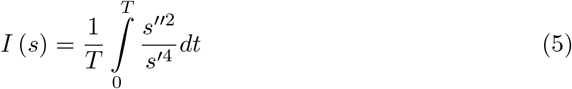

where *T* is the time period required for one cycle of the system; *s*′(*t*) is the tangential speed and *s*″(*t*) is scalar acceleration tangential to the system path in phase space. Both are calculated in terms of the state variables *x_i_* as

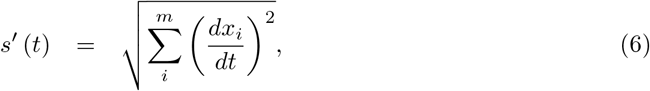

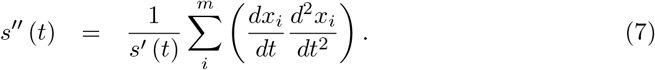

A simple and robust approach to calculating tangential velocity and acceleration uses the three-point difference scheme

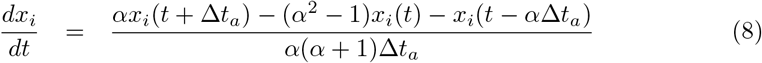

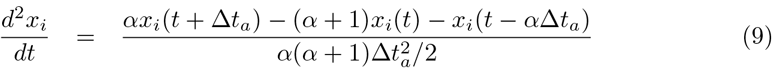

where *x_i_*(*t*) is a central data point, *x_i_*(*t* − Δ*t_a_*) is the later point to the central *x_i_* and *x_i_*(*t* − Δ*t_p_*) is the previous point. For evenly-spaced points Δ*t_a_* = Δ*t_p_* and *α* = Δ*t_p_*/Δ*t_a_* is the ratio of the previous and following time space.

The thesis proposed by [74] is that a change in Fisher information can signal a regime change in a dynamic system, and is based on the following premises: (l) if a change in the dynamic regime is observable then there must be a corresponding change in the measurable variables of the system; (2) an observable change in the measurable variables implies a corresponding change in the distribution of system states; and (3) a change in the distribution of system states implies a change in the system’s Fisher information. An interesting feature of this proposal is that it gives us a way to measure order, since very little information would be inferred from a disordered (non-correlated) system with no observable patterns. This would translate to a Fisher information that approaches zero. On the other hand, the highest values of information are obtained from ordered (highly-correlated) systems that exhibit patterns in behaviour.

Following these ideas, [75] has proposed that:

- Fisher information is a function of the variability of the observations. Low variability leads to high Fisher information and high variability leads to low Fisher information.
- Systems in stable dynamic states have constant Fisher information. Systems losing organization migrate toward higher variability and lose Fisher information.
- Self-organizing systems decrease their variability and acquire Fisher information.

These considerations led them to propose a sustainability hypothesis: “sustainable systems do not lose or gain Fisher information over time.” From equations (5) and (6) this means that the system is in a state of constant tangential velocity and acceleration in the phase space, and therefore, in a stable state.

## Methods

The data were taken from the AmeriFlux researcher-driven network of sites in North, Central and South America, measuring ecosystem respiration, water, and energy fluxes. The network was established to provide compatible data from a large number of sites representing major climate and ecological biomes, including tundra, grasslands, savannah, farmland, and coniferous, deciduous, and tropical forests. Each site has instruments tailored to suit each ecosystem. The network grew from about 15 sites in 1997 to more than 110 active sites registered today. Sixty-one other sites, now inactive, have flux data stored in the network’s database. In 2012, the U.S. DOE established the AmeriFlux Management Project (AMP) at Lawrence Berkeley National Laboratory (LBNL) to support the broad AmeriFlux community and the AmeriFlux sites. The data is publically available from the Ameriflux database (http://ameriflux.lbl.gov). The time series measures CO2 flux fluctuations every half hour.

First, we performed an Analytical Hierarchical Process (AHP) using the following criteria: data requirements for analysis, ecological relevance and quantity of data available. From this analysis, ecosystem respiration was identified as the best measure of ecological processes.

We did not use data from intensively managed farmland sites, because they are subject to high external inputs of nutrients, pesticides and herbicides and are artificially maintained in a temporary ecological condition. In the absence of external control, they would markedly change their physiological signals and so their ecological state should be considered fragile.

In order to derive the Criticality Index and a Scale Invariance Index from annual Ameriflux site data we used the following steps.

Firstly, we extracted the desired variable from the raw file and identified any missing or invalid values in the data series. We then scanned for gaps in the time series, as the computation of the power spectrum demands that the time series have constant time intervals between data values. When the gaps are small (a few data values) it is possible to perform a simple interpolation to fill in the gap. However, to ensure that this interpolation does not alter the data, we only analyzed time series with no gaps at all.

Secondly, we filtered out long-term trends and obvious periodicities from the time series. Although the Ameriflux records are not expected to exhibit long-term trends, we still computed and subtracted a linear trend for the whole time series for each site. If no trend was present the data was left essentially unchanged. Daily periodicities are to be expected for many of the variables. While there are many techniques to extract periodicities for time series, we found that applying a digital infinite impulse response filter worked well for the Ameriflux records. In particular, we used scipy’s notch filter, which is a band-stop filter that rejects a narrow bandwitdh around a chosen frequency and leaves the rest of the spectrum unchanged. We did this three times: once around a frequency of one cycle per day using a quality factor Q=12, and then twice with a narrower band with (Q=30) for the first two harmonics of this frequency. After this step the time series was considered to be trend-free and most high-energy periodicities are removed. We termed this filtered time series the ‘fluctuations’.

The third step was to apply a traditional spectral analysis using a Fast Fourier Transformation of the time series and to compute the spectral index by fitting power-laws to the spectrum. Two fits were obtained. The first is a direct single power-law fit obtained by fitting a straight line to the logarithmic spectrum through least-squares linear regression. The negative of the slope of this line is the first spectral index, *β*_0_, which is a measure of criticality. We used this value to define the Criticality Indicator (Icrit).

The second fit model is a piecewise-defined and double power-law function, composed of a low-frequency power-law with a spectral index *β*_1_, followed by a high-frequency power-law with a spectral index *β*_2_, with a crossover frequency to be determined. We employed the scipy curve-fit routine (a nonlinear least-squares optimizer that uses the Trust Region Reflective algorithm) to obtain the best-fit double power-law model for each of the fluctuation time series. In order to obtain a measure of scale invariance we then compare the two power-law models by computing their Bayesian Information Criterion (BIC) using the residual sum of squares of the models and the relevant number of parameters for each one, including the data variance (i.e. 3 for the single power-law model, 5 for the double-power model). The BIC provides a model comparison that penalizes a model for having more parameters. The BIC values of the models then yields the Scale Invariance Index (Iscale).

We defined the Criticality Index (*Icrit*) as a function of the ‘‘distance” to a 1/*f* type of signal (*β* ~ −1) in such a way that it equals 1 when *β* = −1 and zero when *β* <= −0.5 and *β* >= 1.5. Between those values *Icrit* grows and decreases linearly as *Icrit* = 2*β* − 1 and *Icrit* = −2*β* + 3.

In the same manner, we defined the Scale Invariance Index (*Iscale*) in terms of model selection between a one linear model or a two lines model for fit the PSD using the BIC. Taking the BIC model difference *dBIC* = *BIC*(*model*1) − *BIC*(*model*2), *Iscale* is zero for *dBIC* <= 2 and 1 for *dBIC* > 10. For intermediate values it increases linearly as *Iscale* = (1/8) * *dBIC* − 1.75.

Then, we define the Ecosystemic Health Index (*Ih*) following the functional form of the Human Development Index (HDI) as the square root of the criticality index times the scale index.

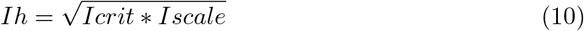

To derive the value of ecosystem stability under the Michaelian framework based on entropy [71], we used the Statcomp library in R [76, 77], which calculates information measures for Time Series, including entropy, complexity and Fisher information, which we report in this work. This library has been used in a range of different disciplinary studies, including medicine [79–86], physical systems [87–91], economic [92–95] and environmental applications [96–98]. Statcomp [76] is based on [78] and calculates simple complexity measures using the concept of permutation entropy defined by the order relations among values of a time series. Permutation entropy assumes that patterns may not have the same probability of occurrence, and that this probability may unveil relevant knowledge about the underlying system. In the same way, we use the *fis* function in the Statcomp library to calculate Fisher information over the annual fluctuation time series.

## Results

In Fig. 3 we show the land condition (*http://www.natureserve.org/conservation–tools/modeling–landscape–condition*) variable used as validation in terms of noise type, see Hak and Comer [100]. The colors represent ecosystem types in the IGBP nomenclature. We expected that if criticality is a good proxy for ecosystem health, then sites with 1/*f* dynamics should have the highest land condition values.

**Fig 3.**
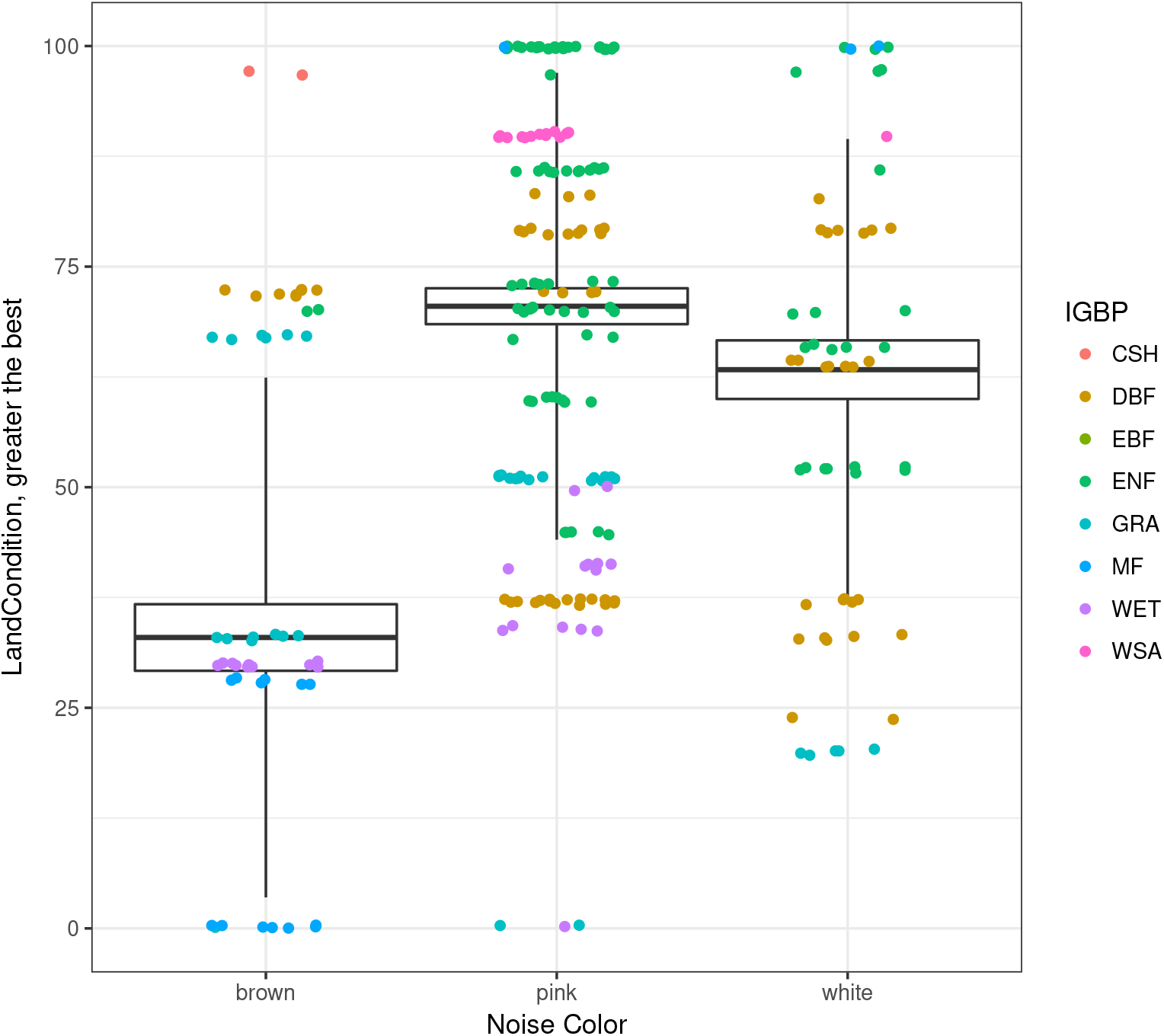
In this figure we show the Land Condition variable used as validation in terms of noise type. Color scale corresponds to ecosystem types in the IGBP nomenclature *gaia.agraria.unitus.it*/*IGBPdesignations.pdf*

By assessing an ANOVA test using land condition as a proxy variable and grouping data by noise type (color) we obtain statistically significant differences (*F* = 28.16; *Pr*(> *F*) = 2.2*x*10^−16^ * **) between noise types, although it is clear that this difference between pink and white might be marginal, perhaps because more data is needed.

Since land condition is not a dynamic measurement, it is not expected to have strong correlation with our measurement of ecosystemic health, but it is consistent in statistical terms. Sites with pink noise (*β* ~ −1) behavior are statistically in better land condition that those sites with white (*β* ~ 0) or brown (*β* ~ −2) noise type. So we may have cases of sites with an external non-health condition that nevertheless exist in a systemically healthy state. An analogy would be a person that has a broken arm: this state of non-health means nothing in terms of systemic processes such as heart activity. Conversely, we also have sites that are externally healthy, but systemically non-healthy. An analogy would be the cases of Sudden Cardiac Death in young athletes [99].

In Fig. 4 we show a boxplot for beta values for each ecosystem type using the IGBPP nomenclature again. As expected, most ecosystems fall into pink noise behavior. And in general, we found that ecosystems out of criticality are older forests, or have been altered by human activity or events such as wildfires. One example is the UA-Me1 (Metolius-Eyerly burn) site, which is an intermediate aged ponderosa pine forest in Oregon-USA that was severely burned in 2002 by the Eyerly wildfire, a stand-replacing event in which all trees were killed.

**Fig 4.**
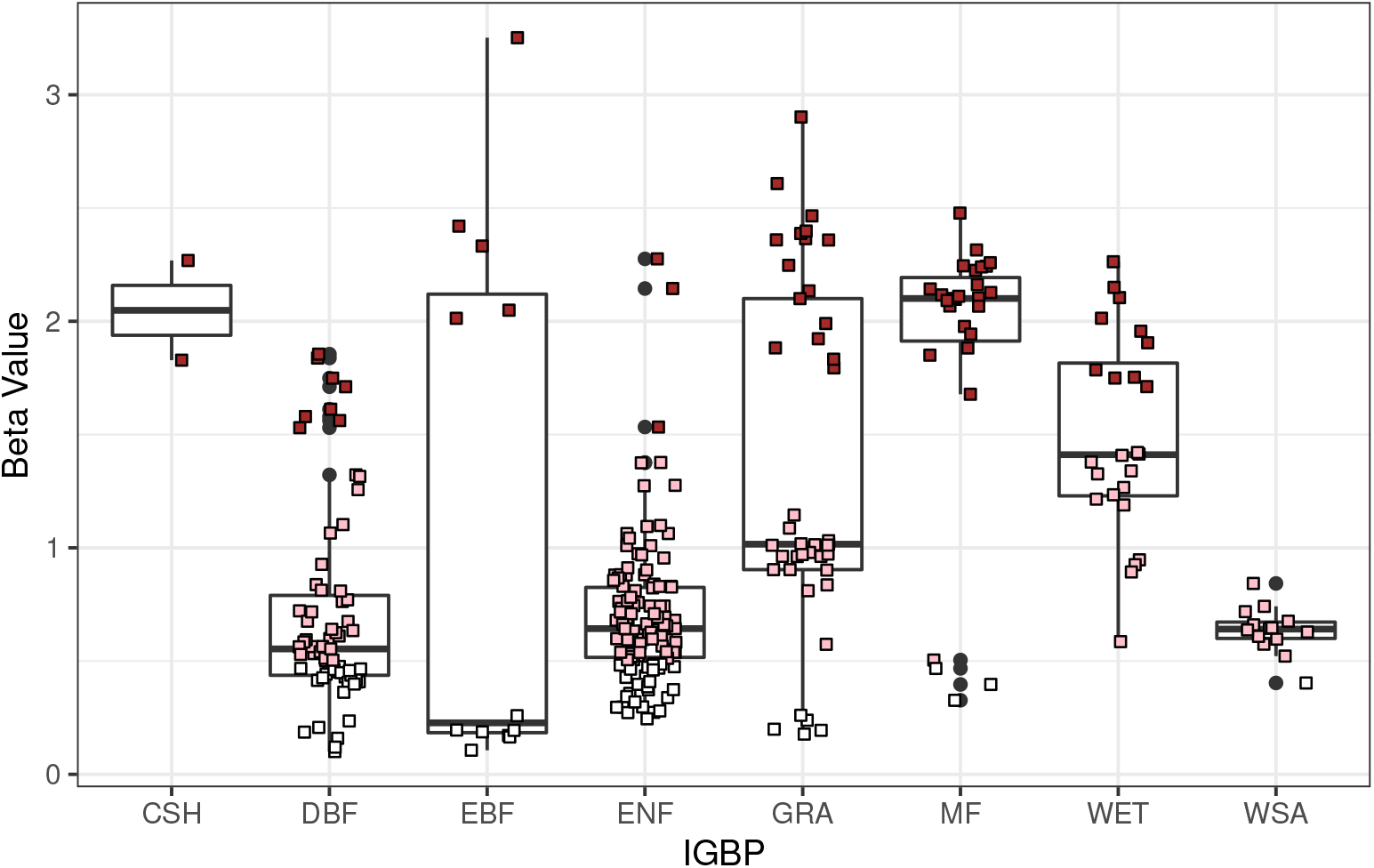
As expected, most ecosystems fall into pink noise. In general, we found that ecosystems out of criticality are older forests or have been altered by human activity or events such as wildfires.

Fig. 5. shows the values of our Ecosystemic Health Index for all ecosystem types (IGBP). For this data set, it seems that Ecosystemic Health is basically driven by the value of beta.

**Fig 5.**
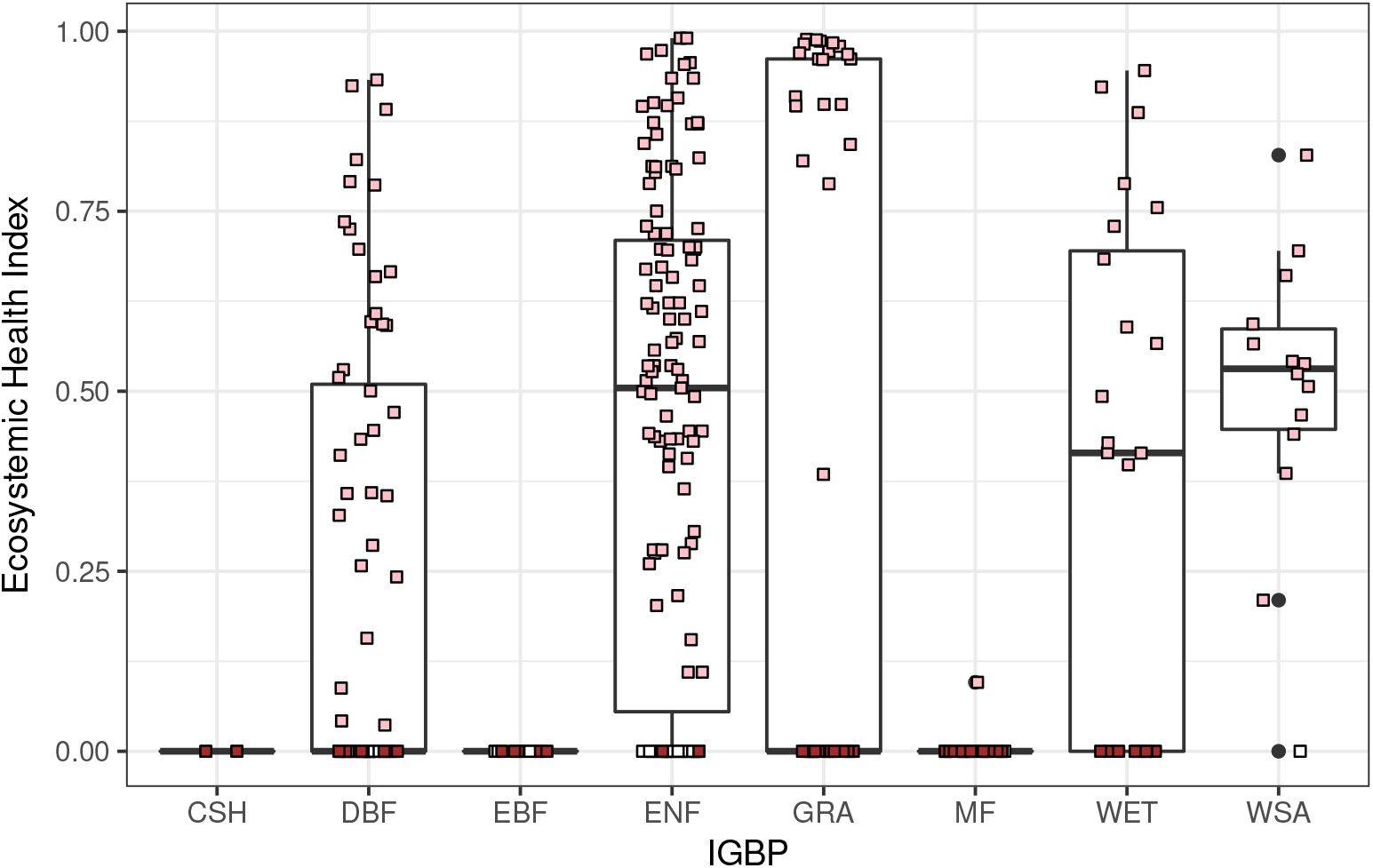
Values of our Ecosystemic Health Index for all ecosystem types (IGBP). The color of each data point corresponds to the type of noise (white, pink and brown)

Our results (see Fig. 6) are also consistent with Michaelian [71] ideas about ecosystem stability and entropy, where stable ecosystems (higher values of entropy) correspond to healthier states (criticality - pink noise).

**Fig 6.**
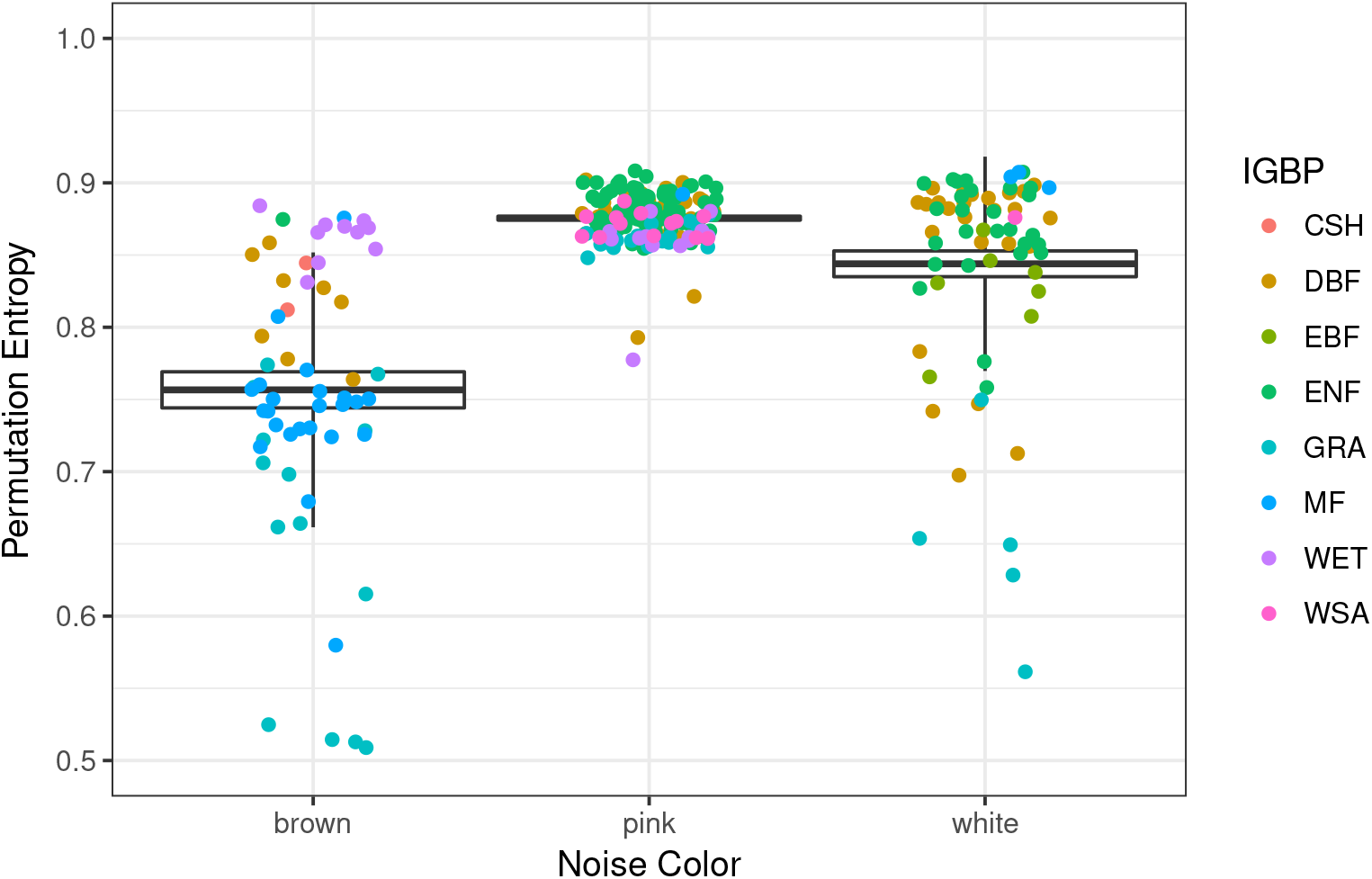
Permutation entropy boxplot in terms of noise color

Again we have significant difference between noise types, with pink noise (*β* ~ −1) behavior being significant higher value of statistical stability than white (*β* ~ 0) or brown (*β* ~ −2) noise.

We see in Fig. 7 a very interesting behavior for permutation entropy as a function of beta. We see that entropy reaches its highest values around the range of beta values for which pink noise is defined, meaning that the most stable behavior corresponds to a criticality dynamic.

**Fig 7.**
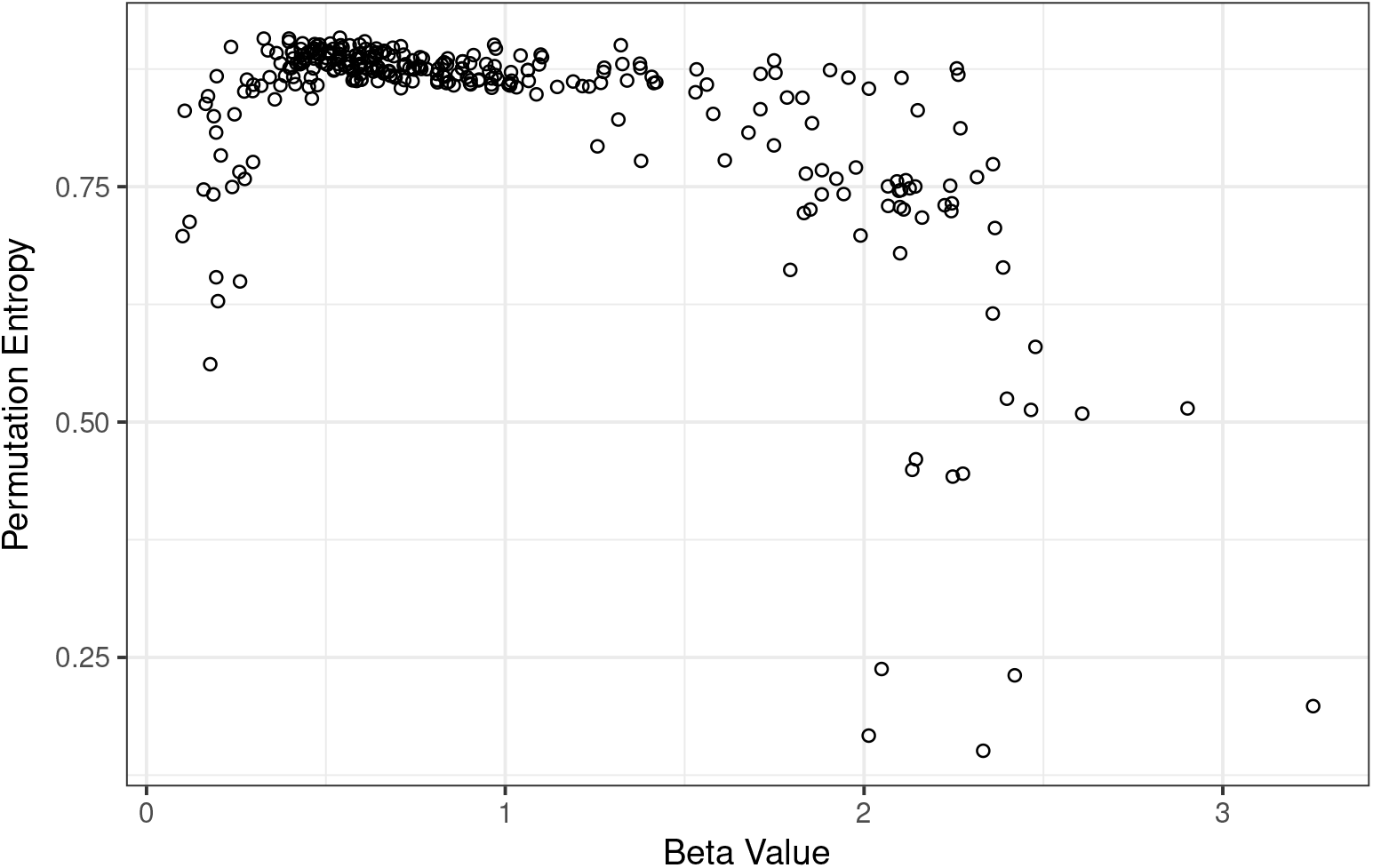
Permutation entropy scatter plot in terms of *β* value. Entropy reaches highest values around the beta values range for which pink noise is defined.

Interestingly enough, as proposed by [21] we see in Fig. 8 a quadratic relation between information (i.e. entropy) and complexity:

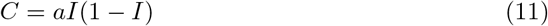

**Fig 8.**
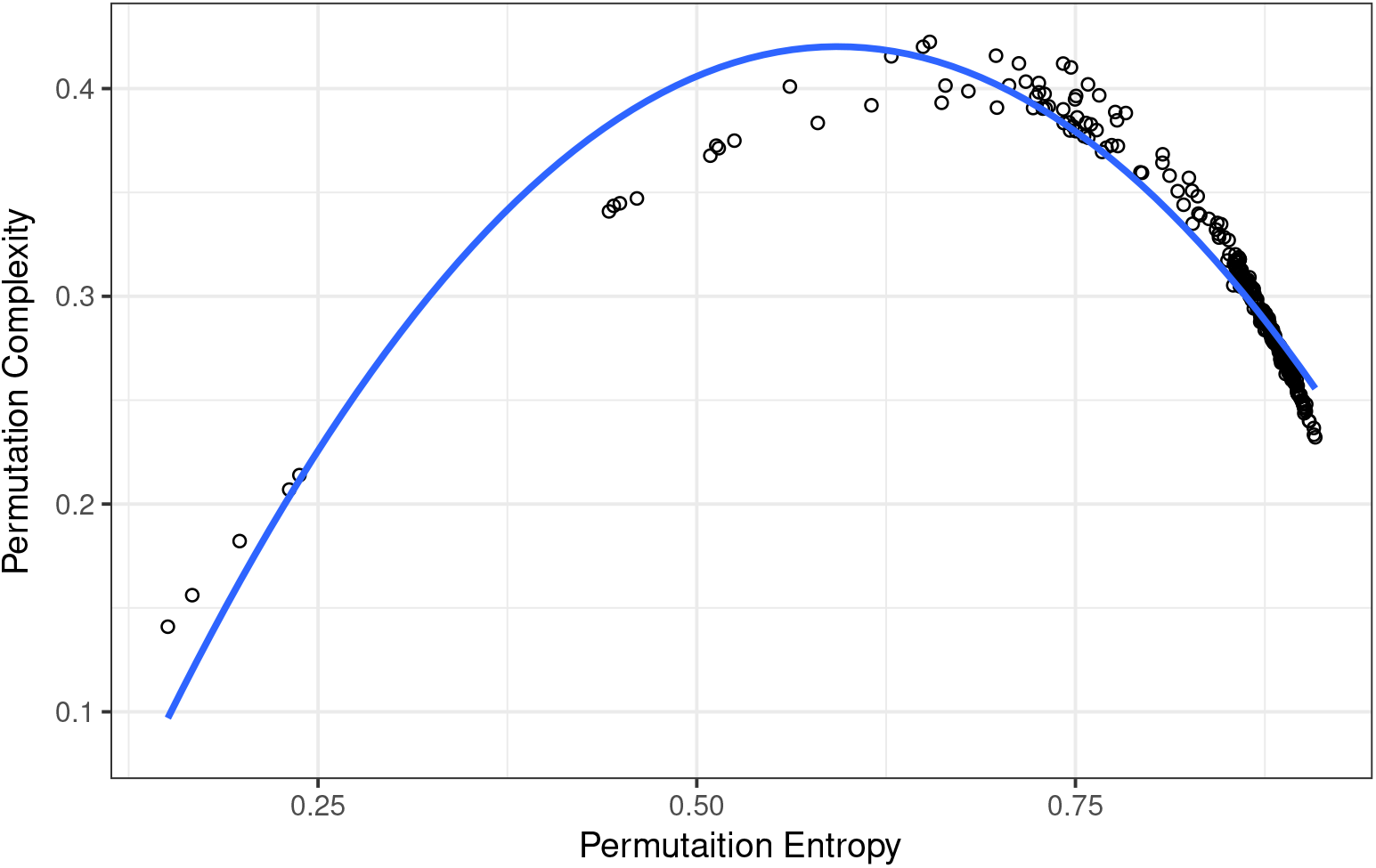
Complexity as a quadratic function of Permutation Entropy.

We think that this might be the first time this relation is found directly in the data and not from modeling.

And finally, our results are consistent with [75]: (a) Fisher information is a function of the variability of the observations such that low variability leads to high Fisher information and high variability leads to low Fisher information; (b) Systems in stable dynamic states have constant Fisher information. Systems losing organization migrate toward higher variability and lose Fisher information; (c) Self-organizing systems decrease their variability and acquire Fisher information. The authors [75] under these considerations propose a sustainability hypothesis: “sustainable systems do not lose or gain Fisher information over time.”

In Fig. 9 we show the evolution of Fisher information for the Harvard Forest site (US-Ha1). We can see that from 1991 to 2003 the ecosystem was in a stable state of low health (combination of white and pink noise) with a low Fisher information value (around 0.025). After that, it enters a process of self-organization gaining Fisher information and starts to stabilize around a higher Fisher information value (around 0.15), dominated by healthier pink noise dynamics, and therefore, according to Cabezas’s hypothesis, a more sustainable state.

**Fig 9.**
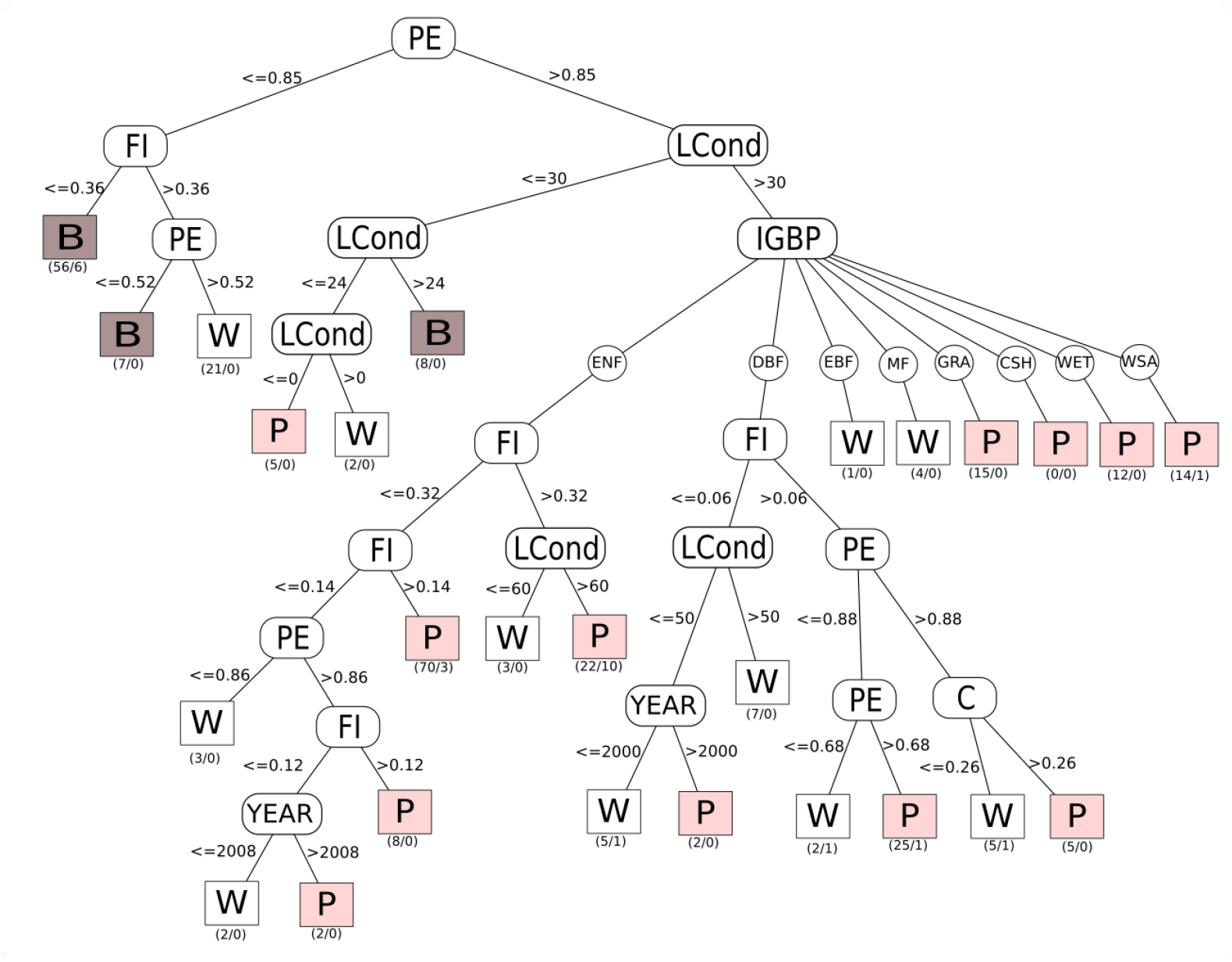
Evolution of Fisher information for the Harvard Forest site (US-Ha1) from 1991 to 2003. Pink points are 1 year time series with a pink noise kind of dynamics (0.5 < *β* < 1.5) and blue point correspond to 1 year time series with a white noise dynamics (*β* ≤ 0.5)

Since criticality (pink noise) appears to be the most healthy and stable (sustainable) type of dynamics, we use it as a leaf variable in a classification tree using the C4.5 algorithm in WEKA (Fig. 10). Results are consistent with what was previously described: sites with an entropy value lower than 0.85 are out of criticality, that is, in non-healthy and non-sustainable states. Sites with a land condition value under 30 are also out of criticality and hence in non-healthy and non-sustainable states. Pink noise in this branch of the tree is a spurious result because, just as with intensive crop lands, this Sherman Island site (US-Snd) is known to be very degraded but under intensive control by the California Department of Water Resources, so this result is a false positive. Specific ecosystem types with a combination of entropy values higher than 0.85 and Land Condition value higher than 30 are in healthy, stable and therefore sustainable states. The rest of the tree is harder to interpret, and individual site histories might play an important role (events of wildfires, site management, and so on.)

**Fig 10.**
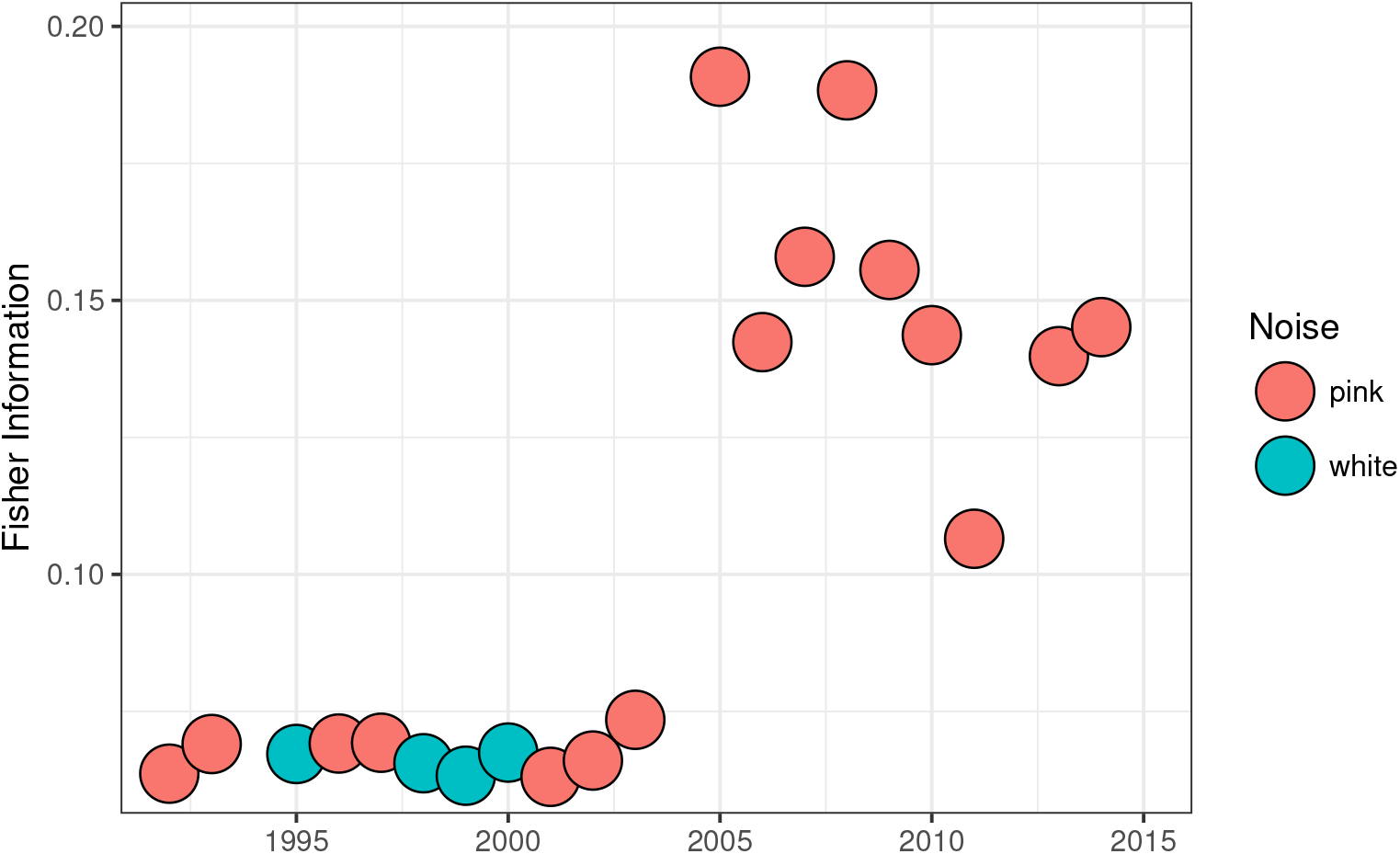
Since criticality (pink noise) appears to be the most healthy and stable (sustainable) type of dynamics, we use it as a leaf variable in a classification tree using the C4.5 algorithm in WEKA.

In Fig. 11 we present the corresponding maps using a combination of circle size and color to encode one or two variables of interest.

**Fig 11.**
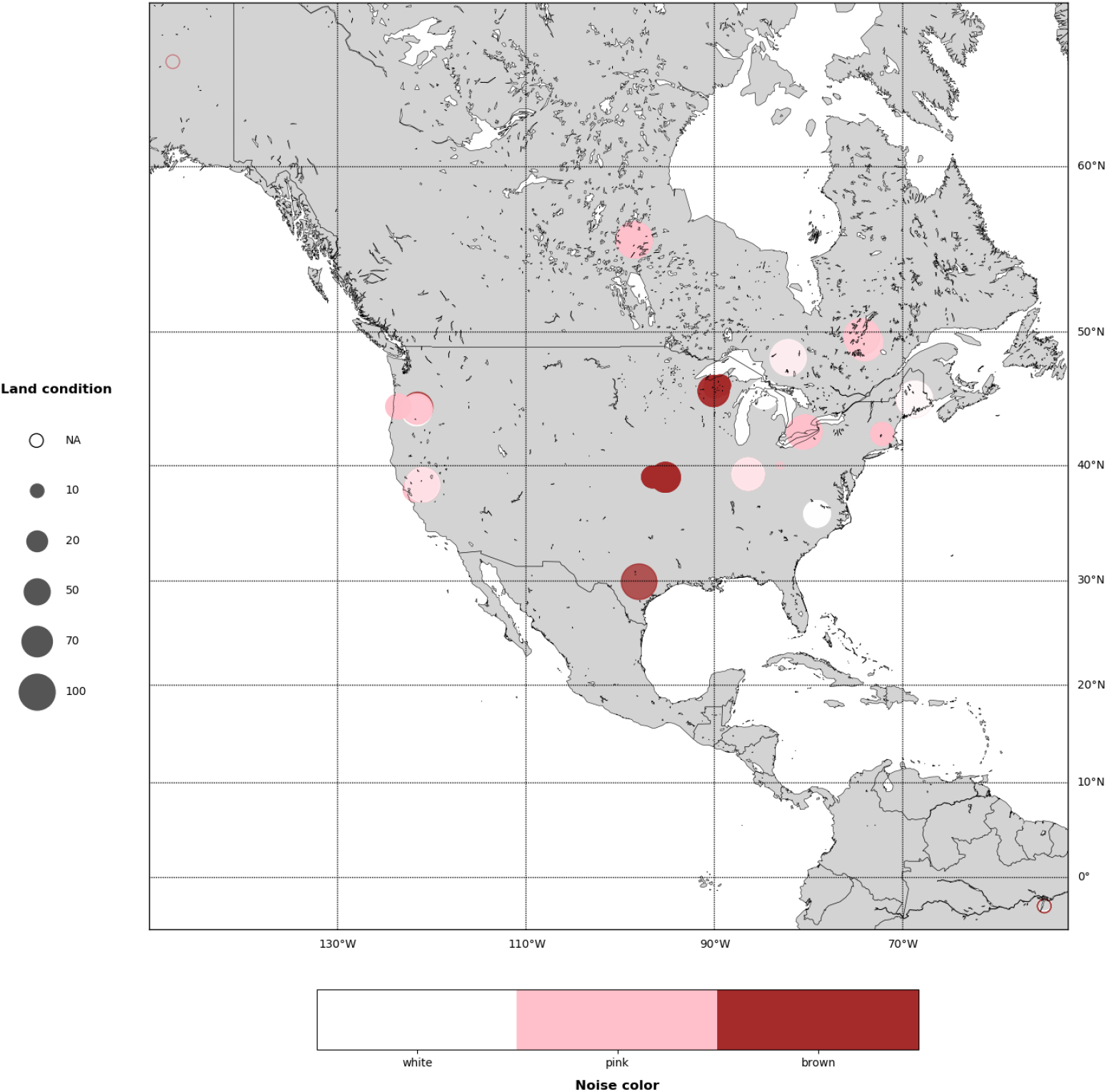
In this map we represent the type of noise in a color scale, and land condition as size of the circles.

Finally we show the complete Sustainability Index as the square root of the Health Index times the Stability Index Fig. 12.

**Fig 12.**
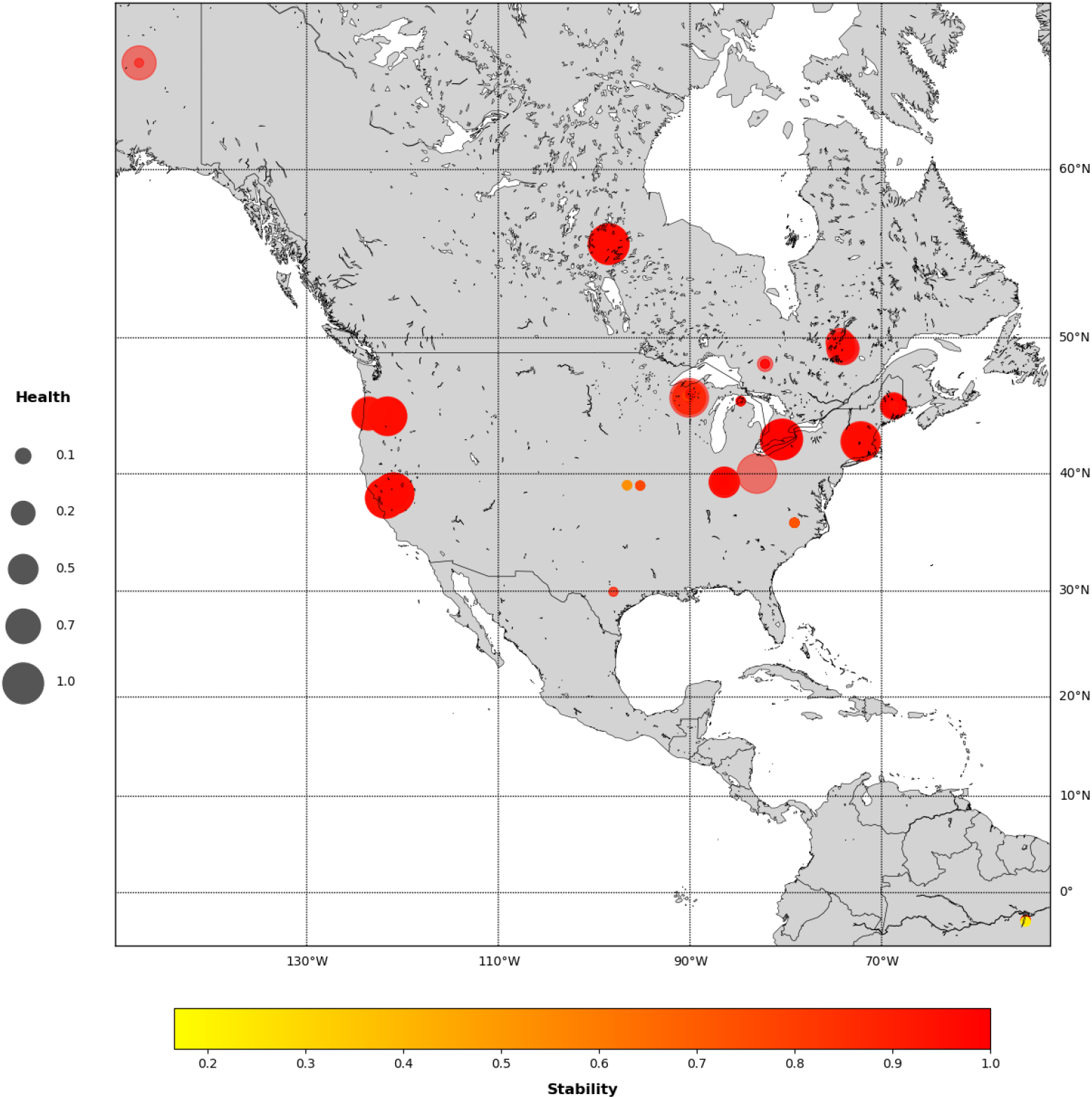
Here, Ecosystem Stability (Permutation Entropy) is shown in a color scale and Ecosystem Health (as the square root of scale invariance indicator times the criticality indicator) as the size of the circles.

## Discussion and conclusions

A complexity perspective based on information theory seems to be a promising starting point to develop a general framework for the measurement of sustainability.

We showed that the use of criticality, defined by scale invariance and pink noise behavior, may be one way to measure systemic ecosystem health. We complement the analysis using a non-dynamic variable, in this case land condition. In general, we found that sites with pink noise (*β* ~ −1) behavior are statistically in better land condition that those sites with white (*β* ~ 0) or brown (*β* ~ −2) noise type. Interestingly enough, one may find systems with good land condition but in low systemic health and vice versa.

We interpret the first case, where we have high values of land condition but low values of Ecosystem Health, in terms of an analogy with human health considering phenomena such as Sudden Cardiac Death in young athletes [99]. Consider an Olympic athlete in their twenties: it would be difficult to think of someone with better external health qualities, and yet this athlete could drop dead on the track due to this syndrome, which is related to systemic health. On the other hand, consider a person with a broken arm (a clear external signal of non-health) but perfectly healthy in terms of heart, brain and general system functioning. Additionally, we identified a third case for intensively managed ecosystems that, as in this analogy, matches a patient in intensive care who is maintained in some form of artificial health using life-support devices.

We recognize that as usually happens with complex systems: the basis of the configuration space of the problem is not known a priori. Whether ecosystem respiration is the correct or only “physiological” signal for Ecosystemic Health measurements remains an open question.

Our results are consistent with stability ideas developed by Michaleian, where high entropic sites are also in criticality. We refer to a case study where our results are also in good agreement with the Fisher information framework for system stability developed by Mayer and co-workers [3]. In this study, stable (more sustainable) systems do not lose or gain Fisher information over time.

For future work, other sources of time series should be systematically explored, for example data already available in public repositories such as:

- http://dataportal-senckenberg.de/knb/
- https://www.st.nmfs.noaa.gov/copepod/time-series/
- https://eco.confex.com/eco/2017/webprogram/Paper62280.html
- https://lagoslakes.org/

In particular we think that time series from main biogeochemical processes could be good candidates for systemic ecosystem physiological signals. For example, biologically available nitrogen (fixed N) limits the fertility of much of the ocean and data is available in the NOAA link above.

Finally, we acknowledge that more data and controlled experiments would be necessary to understand complicated patterns that emerge from current analysis.

